# Strong Purifying Selection Contributes to Genome Streamlining in Epipelagic *Marinimicrobia*

**DOI:** 10.1101/653261

**Authors:** Carolina Alejandra Martinez-Gutierrez, Frank O. Aylward

**Affiliations:** Department of Biological Sciences, Virginia Tech, Blacksburg, VA

**Keywords:** genome streamlining, purifying selection, evolutionary genomics, dN/dS ratio, *Marinimicrobia*, marine bacterioplankton

## Abstract

Marine microorganisms inhabiting nutrient-depleted waters play critical roles in global biogeochemical cycles due to their abundance and broad distribution. Many of these microbes share similar genomic features including small genome size, low % G+C content, short intergenic regions, and low nitrogen content in encoded amino acids, but the evolutionary drivers of these characteristics are unclear. Here we compared the strength of purifying selection across the *Marinimicrobia*, a candidate phylum which encompasses a broad range of phylogenetic groups with disparate genomic features, by estimating the ratio of non-synonymous and synonymous substitutions (dN/dS) in conserved marker genes. Our analysis shows significantly lower dN/dS values in epipelagic *Marinimicrobia* that exhibit features consistent with genome streamlining when compared to their mesopelagic counterparts. We found a significant positive correlation between median dN/dS values and genomic traits associated to streamlined organisms, including % G+C content, genome size, and intergenic region length. Our findings are consistent with genome streamlining theory, which postulates that small, compact genomes with low G+C contents are adaptive and the product of strong purifying selection. Our results are also in agreement with previous findings that genome streamlining is common in epipelagic waters, suggesting that genomes inhabiting this region of the ocean have been shaped by strong selection together with prevalent nutritional constraints characteristic of this environment.

## Main Text

Bacteria and Archaea play key roles in marine biogeochemical cycles and are a dominant force that drives global nutrient transformations (Azam et al. 1983; Falkowski et al. 2008). Our understanding of microbial diversity in the ocean has been transformed in the last decades due to the discovery of several globally-abundant marine microbial lineages that are among the most numerically abundant life forms on Earth (Giovannoni & Stingl 2005). Work on some of these abundant lineages succeeded in culturing representatives that could then be studied extensively in the laboratory, such as *Prochlorococcus marinus* (Chisholm et al. 1992) and heterotrophic bacterioplankton belonging to the *Pelagibacteriales* (Rappé et al. 2002) and *Roseobacter* groups (Luo & Moran 2014), but many other dominant microbial lineages have not been brought into pure culture and require cultivation-independent methods for analysis (DeLong & Karl 2005).

Previous research of *Prochlorococcus* and *Pelagibacter* genomes provided some of the earliest insights into the ecology and evolution of these dominant planktonic microbial lineages (Partensky & Garczarek 2010; Giovannoni 2005; Giovannoni et al. 2014). It was quickly noted that both groups had small genomes that contained short intergenic regions and encoded among the fewest genes of any free-living organism (Giovannoni et al. 2014). These characteristics were explained through the proposed theory of genome streamlining, which states that genome simplification is an adaptation to consistently oligotrophic conditions, and therefore the loss of non-coding DNA and unnecessary genes (and corresponding transcriptional, translational, and regulatory burdens) is beneficial (Giovannoni et al. 2014). Genome streamlining theory is supported by the observation that many streamlined genomes also have a higher proportion of Adenine and Thymine nucleotides (i.e., low %GC content) and their encoded proteins are correspondingly enriched in amino acids that have a low nitrogen content, which would be expected to be beneficial in oligotrophic waters (Giovannoni et al. 2014; Grzymski & Dussaq 2012). More recent cultivation-independent studies of marine lineages have confirmed that genomic features consistent with genome streamlining are prevalent in a variety of marine lineages in addition to *Prochlorococcus* and *Pelagibacter* (Ghai et al. 2013; Swan et al. 2013; Luo et al. 2014; Getz et al. 2018; Dupont et al. 2012), suggesting that common evolutionary drivers shape diverse bacterioplankton groups in the ocean.

Although the term “genome streamlining” implies adaptation under oligotrophic nutrient conditions, it remains a possibility that these genomic signatures are non-adaptive and instead the result of processes other than strong purifying selection (Batut et al. 2014). For example, it has long been known that endosymbiotic bacteria contain small genomes with short intergenic regions and low % GC content, but in these cases the drivers are a small effective population size (*N*_*e*_) and consequently high genetic drift, which allows slightly deleterious deletions to become fixed (Kuo et al. 2009; Charlesworth 2009; McCutcheon & Moran 2011). While it remains unlikely that marine free-living bacteria have small effective population sizes comparable to those of endosymbiotic bacteria (Biller et al. 2015), it has been argued that population bottlenecks in the distant evolutionary past of some marine lineages may have precipitated their initial gene losses (Luo et al. 2017). Moreover, recent work has also shown that weakly deleterious mutations and low recombination rates can substantially lower the efficacy of purifying selection in bacterial genomes (Price & Arkin 2015), implying that the large abundances of marine bacteria may not translate directly into high selection.

In this study we sought to test whether streamlined marine genomes experience higher levels of purifying selection than non-streamlined genomes. This is a key prediction of genome streamlining theory that, if correct, would strongly suggest that genomic features associated with genome streamlining are adaptive and not a result of genetic drift. We focused our analyses on *Marinimicrobia*, a diverse marine phylum that comprises globally-abundant lineages involved in distinct biogeochemical processes (Hawley et al. 2017; Getz et al. 2018). The *Marinimicrobia* are an ideal group to test genome streamlining theory because genomic traits vary widely across this phylum together with its distribution in the water column, with features consistent with genome streamlining evolving multiple times independently in different clades (Getz et al. 2018). We would expect that *Marinimicrobia* that live in epipelagic waters and tend to exhibit features of genome streamlining, such as low % GC content, short intergenic spacers, and relatively low nitrogen content but carbon-rich encoded amino acids, will therefore show higher levels of purifying selection than mesopelagic *Marinimicrobia*, which generally have opposing genomic characteristics (Getz et al. 2018).

In order to test our hypothesis, we estimated the ratio of nonsynonymous and synonymous substitutions (dN/dS) of highly conserved marker genes. In general, dN/dS values <1 are indicative of purifying selection, thus the relative selection strength can be compared across groups using this metric since lower values imply higher levels of selection strength (Kryazhimskiy & Plotkin 2008). To ensure that our results could be accurately compared across different clades, we used two sets of marker genes that are broadly shared among Bacteria, which we refer here to as the EMBL and CheckM marker gene sets (Sunagawa et al. 2013; Parks et al. 2015). Our results showed that median dN/dS *Marinimicrobia* values span two orders of magnitude for the CheckM data set, between 0.1912 and 0.001, and one order of magnitude when analyzing the EMBL marker genes, between 0.2383 and 0.0161 (Fig. 1). The values obtained are far lower than one, which is consistent with the expectation that conserved phylogenetic marker genes experience purifying selection in order to maintain protein function (Novichkov et al. 2009). We analyzed the dN/dS values in the context of the *Marinimicrobia* phylogeny and identified a general pattern in which epipelagic genomes exhibiting streamlined features showed lower median dN/dS values than non-streamlined mesopelagic genomes (Fig. 1). This differentiation was particularly evident within clade 2, which encompasses epipelagic and mesopelagic organisms within the same branch (Fig. 1, in red). The observation of lower dN/dS values in epipelagic *Marinimicrobia* was strongly supported by statistical analyses of both the CheckM and the EMBL marker gene sets (P<0.0001 and P<0.0001, respectively; Fig. 2). Our findings suggest that *Marinimicrobia* found in different habitats exhibit distinct selection strength, with epipelagic *Marinimicrobia* experiencing stronger purifying selection.

**Fig. 1:**
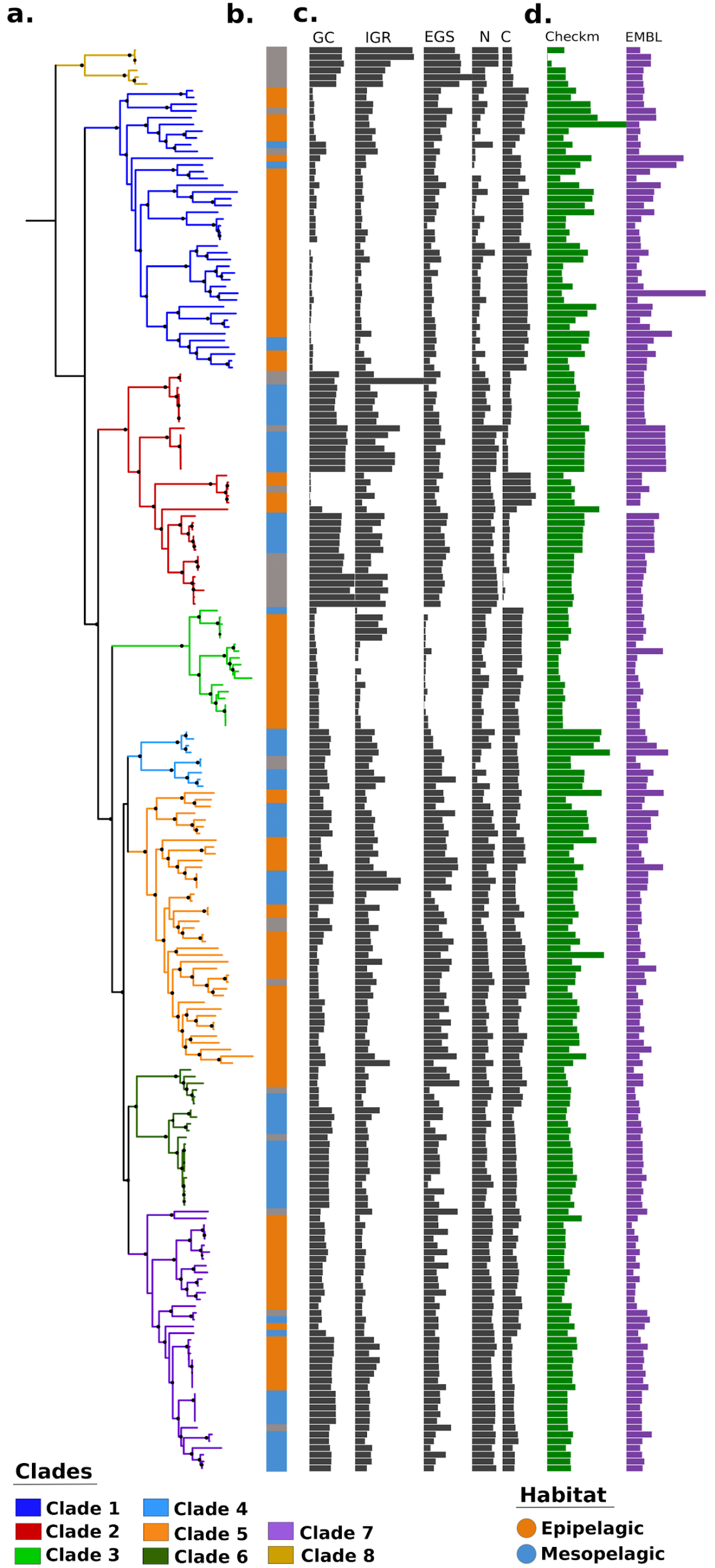
Representation of phylogeny, habitat classification, genomic features, and median dN/dS values of the *Marinimicrobia* phylum. (a) Phylogenetic tree of the 211 genomes constructed using amino acids sequences of 120 highly conserved marker genes. (b) Habitat classification of *Marinimicrobia* genomes based on Getz et al. (2018), grey bars represent unknown habitat. c) Genomic features of the Marinimicrobial genomes. Abbreviations: GC, % GC content (range, 27 to 55%); IGR, mean intergenic region length (range, 30 to 187 nucleotides); EGS, estimated genome size (range, 1 to 4.3 Mbp); N-ARSC (range, 0.3 to 0.34); C-ARSC (range, 3 to 3.2). (d) Median dN/dS values calculated based on two marker genes data sets. Checkm range between 0.001 and 0.1912; EMBL range between 0.0161 and 0.2383. Black points on branches represent support values of >95%.

**Fig. 2:**
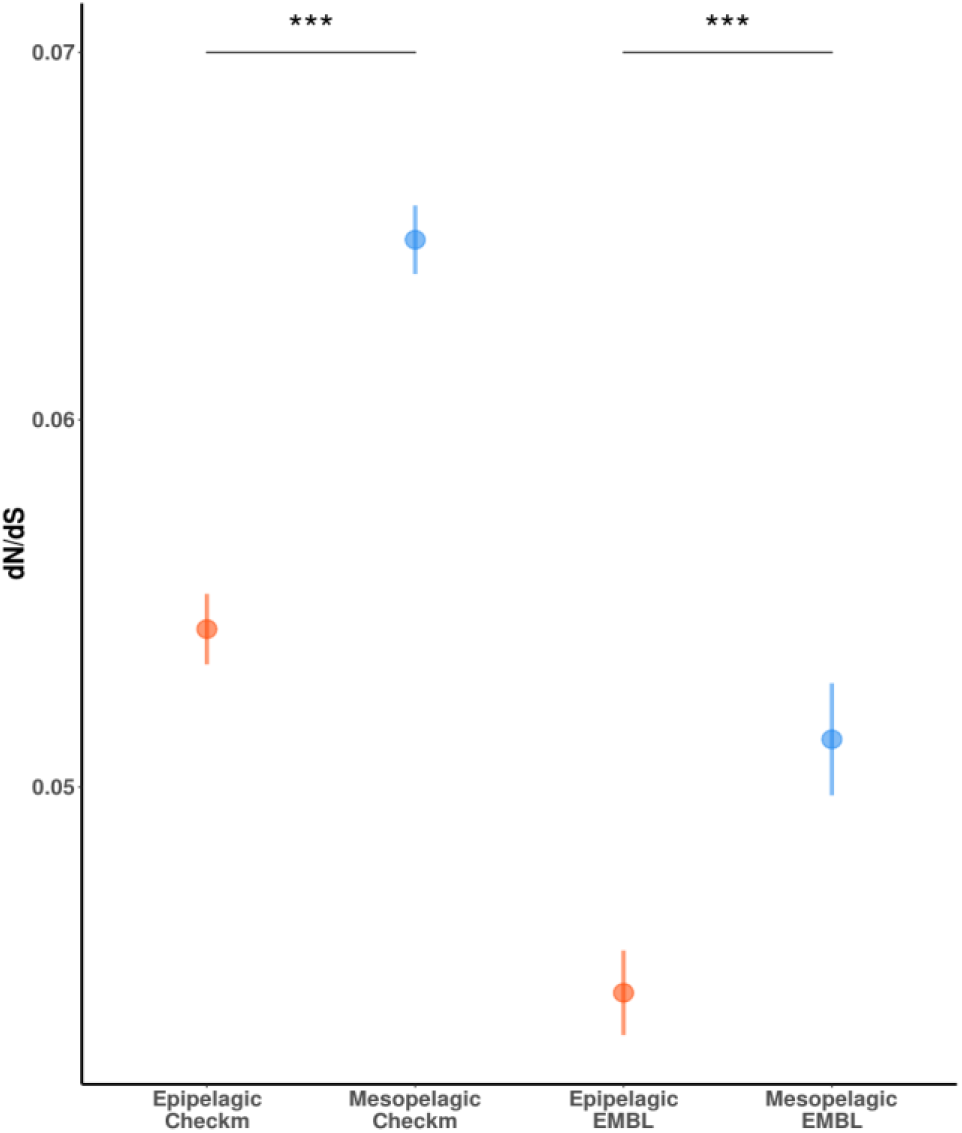
Plot representing median dN/dS values of epipelagic and mesopelagic *Marinimicrobia.* Bars show standard error. Statistical significance of differences between dN/dS values groups according to a non-paired, one-sided Mann Whitney-Wilcoxon test is denoted by: (***) for *P*<0.0001.

We additionally explored the correlation between the strength of selection represented by the dN/dS ratio and several genomic features associated with streamlining. It was found that genomes showing the lowest % GC content have the lowest dN/dS values (Spearman’s rho = 0.41, P<0.0001; Fig. 3a), and thus the higher selection strength. The same trend was obtained when correlating the dN/dS ratio with the estimated genome size (Spearman’s rho = 0.29, P= 0.0002; Fig. 3b) and intergenic spacers length (Spearman’s rho = 0.52, P<0.0001; Fig. 3c). To analyze the carbon and nitrogen content of encoded proteins we estimated the carbon and nitrogen atoms per residue side chain (C-ARSC and N-ARSC, respectively), which have previously been used for this purpose (Grzymski & Dussaq 2012; Getz et al. 2018; Mende et al. 2017). The correlation between C-ARSC and dN/dS showed a negative relationship (Spearman’s rho = −0.30, P< 0.0001; Fig. 3d), whereas N-ARSC correlation showed an increase along dN/dS values (Spearman’s rho = 0.26, P= 0.0005; Fig. 3e). This observation is consistent with previous findings that carbon-rich proteins with respect to nitrogen are prevalent in surface waters and suggests that this feature may be advantageous under the conditions found in this environment (Mende et al. 2017).

**Fig. 3:**
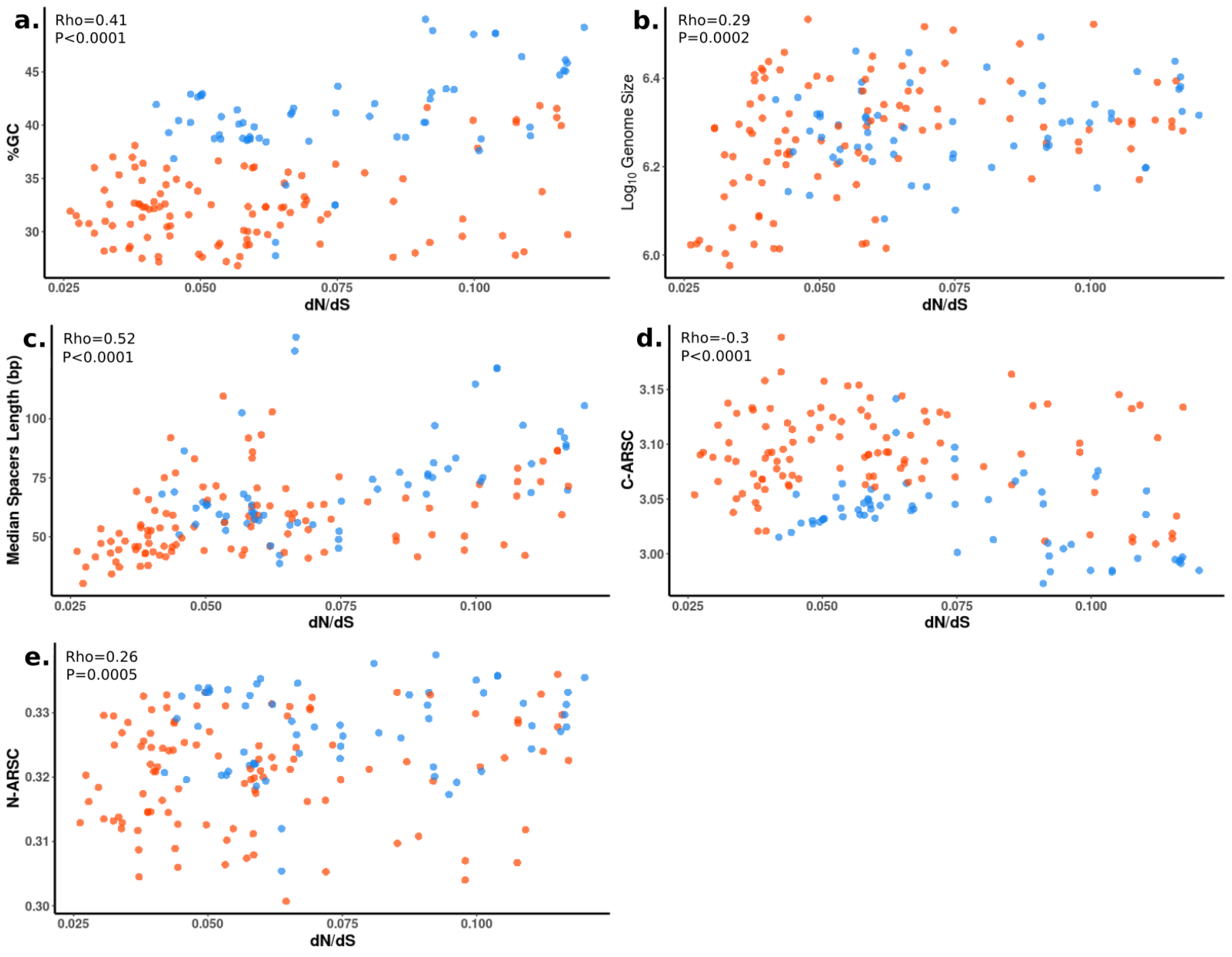
Scatter plots showing the relationship between median dN/dS values and streamlined genomic features of the *Marinimicrobia* genomes. Median dN/dS values were calculated including both data sets. Orange and blue points denote epipelagic and mesopelagic genomes, respectively. (a) Median dN/dS vs. %GC content. (b) Median dN/dS vs. Log_10_ estimated genome size. (c) Median dN/dS value vs. median spacers length. (d) Median dN/dS vs. C-ARSC. Spearman correlation was performed for each variables pair and details can be consulted on the main text.

Because we evaluated multiple genomic features that have been identified as indicators of streamlining (Giovannoni 2005; Giovannoni et al. 2014), we performed a multivariate analysis to explore the effect of such genomic variables on streamlined and non-streamlined genomes differentiation. The first two main axes obtained from our principal component analysis (PCA) explained 71% of the variance (Fig. 4a). Additionally, we observed a tight clustering of *Marinimicrobia* genomes when environment was included as variable (Fig. 4b). These findings suggest that the strength of selection is consistently different between epipelagic and mesopelagic *Marinimicrobia*, and that strong purifying selection is correlated with genomic characteristics associated with genome streamlining. It is important to note that these results do not imply that strong selection will always lead to low % GC content or high C-ARSC, however, since the genomic changes that result from strong selection depend on prevailing environmental factors. For example, strong selection in mesopelagic waters would not necessarily lead to an increase in amino acid carbon content since carbon is relatively more limiting in deep waters compared to other macronutrients. The disparate genomic features of epipelagic and mesopelagic *Marinimicrobia* are therefore likely the result of differential nutrient availability along the water column; surfaces waters have long been characterized as nutrient-depleted (i.e., nitrogen and iron) but carbon-rich due to photosynthetic activities (Grzymski & Dussaq 2012; Mende et al. 2017) in comparison to the mesopelagic zone, where photosynthesis is limited but particulated organic matter remineralization occurs, thus nitrogen is relatively more abundant (Karl 2002; Moore et al. 2013; Grzymski & Dussaq 2012).

**Fig. 4:**
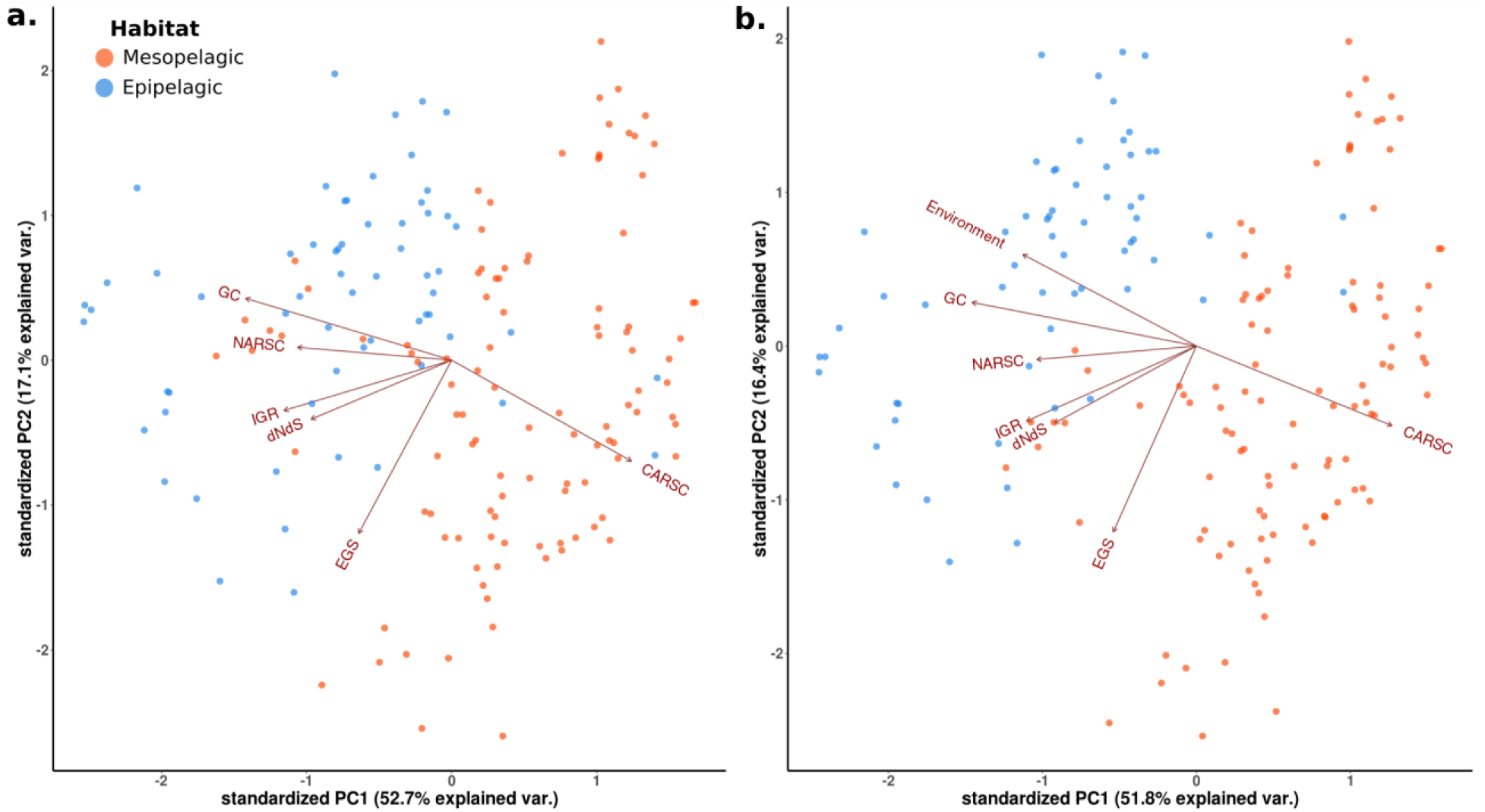
PCA analysis displaying the Euclidean distance among Marinimicrobial genomes. (a) PCA excluding Environment as variable. (b) PCA including Environment as variable. Abbreviations: GC, %GC content; NARSC, N-ARSC; IGR, mean intergenic region length; dNdS, median dN/dS ratio based on both marker gene data sets; EGS, estimated genome size; CARSC, C-ARSC.

An important caveat of dN/dS ratio is that it only provides insight into the strength of recent selective pressure and therefore cannot be used to infer the selective strength experienced by lineages in the distant past. Other streamlined lineages such as the *Pelagibacterales* and *Prochlorococcus* are thought to have underwent genome reduction in the distant past, and it is therefore difficult to assess the strength of selection on these ancestral genomes during these transitions. Some studies have suggested that genetic drift due to possible population bottlenecks drove these genomic changes (Luo et al. 2017), while other studies have suggested strong purifying selection was the primary driver (Sun & Blanchard 2014). In contrast, *Marinimicrobia* appear to have experienced multiple independent genome transition events more recently in their evolutionary history (Getz et al. 2018), and comparison of the selective pressures across disparate clades with similar genomic features therefore provides insight into the selective regimes that led to current genomic architectures. Our finding of consistently higher purifying selection in streamlined epipelagic *Marinimicrobia* from multiple different clades therefore suggests that streamlining is an adaptation and therefore not a product of high genetic drift.

## Materials and Methods

### *Marinimicrobia* genomes collection

We analyzed a set of 211 *Marinimicrobia* genomes that was previously compiled (Getz et al. 2018). This data set included all genomes in GenBank classified as *Marinimicrobia* according to the NCBI Taxonomy available until 15 October 2017 (Sayers et al. 2019), as well as *Marinimicrobia* genomes from the Integrated Genomes system (IMG (Markowitz & Kyrpides 2007)), and from two different studies in which metagenome-assembled genomes (MAGs) were generated (Tully et al. 2018; Delmont et al. 2018). We quality-filtered the genomes based on the results of CheckM (Parks et al. 2015), with only those genomes with contamination levels of <5% and completeness of >40% were considered for further analyses. Additionally, in order to remove genomes incorrectly classified and redundant, a second filtering step was performed based on a preliminary multilocus phylogenetic tree (Getz et al. 2018). The data set employed by Getz *et al.* was complemented with the genomes SCGC_AD-604-D17, SCGC_AD-606-A07, SCGC_AD-615_E22, TOBG_RS-419 (Tully et al. 2018; Delmont et al. 2018).

### Phylogenetic reconstruction

To reconstruct the *Marinimicrobia* phylogeny, we predicted proteins from genomes using Prodigal v2.6.2 (Hyatt et al. 2010) and identified phylogenetic marker genes using HMMER3 (Eddy 2011). We constructed a phylogeny from an amino acid alignment created from the concatenation of 120 marker genes that have been previously used for phylogenetic reconstruction of Bacteria (Parks et al. 2015). The trusted cutoffs were used in all HMMER3 searches with the “cut_tc” option in hmmsearch. We used the standard_fasttree workflow included in the ETE Toolkit which includes ClustalOmega for alignment (Sievers & Higgins 2018), trimAl for alignment trimming (Capella-Gutierrez et al. 2009), and FastTree for phylogenetic estimation (Price et al. 2010). The different branches obtained were classified into clades based on previously published results (Getz et al. 2018). We visualized the resulting tree as well as genomic features and median dN/dS values for each genome in the interactive Tree of Life (iTOL(Letunic & Bork 2016); https://itol.embl.de/tree/45372154390391554311793).

### dN/dS ratio calculation and filtering

To estimate the strength of purifying selection we used the ratio of nonsynonymous and synonymous substitutions (dN/dS) ratio, which has been widely used for this purpose. Although the absolute value of the dN/dS ratio will vary depending on the gene used, in general lower values are a sign of higher purifying selection while higher values are a sign of higher genetic drift (low purifying selection). To calculate genome-wide dN/dS ratios we used two sets of conserved marker genes that would be expected to be found in most genomes. The first one consists of 120 phylogenetic marker genes that is highly conserved in Bacteria, which we refer to as the Checkm set due to its use in the CheckM tool (Parks et al. 2015). The second set consists of 40 phylogenetic marker genes that has been used in inter-domain phylogenetics, which we refer to as the EMBL set due to its development in several papers published at the European Molecular Biology Laboratory (Sunagawa et al. 2013).

For both marker gene sets, we predicted proteins from each genome using Prodigal and then annotated the marker genes of interest using the hmmsearch tool of HMMER3 with the recommended cutoffs (Eddy 2011). We aligned the amino acid sequences for each annotated gene coming from *Marinimicrobia* genomes separately using ClustalOmega (Sievers & Higgins 2018), and the resulting alignments converted into codon alignments using PAL2NAL (Suyama et al. 2006). Maximum-likelihood approximation (codeML) within the PAML 4.9h package (Yang 2007) was used through Biopython in order to perform dN/dS pairwise comparison within the clades previously established (Getz et al. 2018). We removed dN/dS values with dS > 2 and dS < 0.01, which implies saturation of synonymous substitutions and highly dissimilar sequences that provide unreliable estimates, respectively (Ran et al. 2014). Additionally, we discarded all dN/dS values > 10 on the grounds that these were largely artefactual.

### Calculation of genomic features associated to streamlining

We estimated GC content, intergenic regions length, C-ARSC, and N-ARSC through code previously developed (Mende et al. 2017). For genome length estimation, the following equation was used:

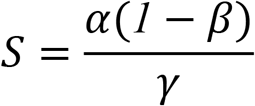

Where α is the number of base pairs in the genome assembly, β is the estimated level of contamination, and γ is the estimated level of completeness. Contamination and completeness for each genome were determined using CheckM (Getz et al. 2018).

### *Marinimicrobia* genomes distribution and statistical analyses

The ratio of non-synonymous and synonymous substitutions is a widely used metric that estimates the evolutionary pressure on protein-coding genes (Kryazhimskiy & Plotkin 2008). In order to investigate the strength of selection acting on epipelagic and mesopelagic *Marinimicrobia*, genomes were classified into epipelagic and mesopelagic based on their biogeographic distribution (Getz et al. 2018). For statistical analysis, we loaded the filtered dN/dS for both data sets and each genome into R and performed mean comparisons analysis through Mann-Whitney U test using the “wilcox.test” function. Additionally, in order to investigate the relationship between median dN/dS and genomic features associated to each genome, we applied the “cor.test” function with the Spearman method. Mean comparisons and correlation plots were visualized through the ggplot2 package (Wilkinson 2011). Also, we explored the distance between epipelagic and mesopelagic *Marinimicrobia* genomes employing the genomic features and median dN/dS values through a PCA analysis with the “prcomp” function available on R. Euclidean distance was visualized using the “ggbiplot” function within the ggplot2 package (Wilkinson 2011).

## Acknowledgments

We acknowledge use of the Virginia Tech Advanced Research Computing Center for bioinformatic analyses performed in this study. This work was supported by the Institute for Critical Technology and Applied Science at Virginia Tech and a Sloan Research Fellowship and a Simons Early Career Award in Marine Microbial Ecology and Evolution to FOA.

